# Identifying Optimal Neuroinflammation Treatment Using Nanoligomer™ Discovery Engine

**DOI:** 10.1101/2022.08.23.505002

**Authors:** Sadhana Sharma, Curtis Borski, Jessica Hanson, Micklaus A. Garcia, Christopher D. Link, Charles Hoeffer, Anushree Chatterjee, Prashant Nagpal

**Affiliations:** Sachi Bioworks, 685 S Arthur Avenue, Colorado Technology Center, Louisville, CO 80027; Department of Integrative Physiology, University of Colorado Boulder, Boulder, CO 80309; Institute for Behavioral Genetics, University of Colorado Boulder, Boulder, CO 80303

**Keywords:** Chronic neuroinflammation, LPS-induced neuroinflammation, gene therapy, drug discovery, target validation, neurodegeneration, sleep countermeasure

## Abstract

Acute activation of innate immune response in the brain, or neuroinflammation, protects this vital organ from a range of external pathogens and promotes healing after traumatic brain injury. However, chronic neuroinflammation leads to the activation of immune cells like microglia and astrocytes causes damage to the nervous tissue, and is causally linked to a range of neurodegenerative diseases such as Alzheimer’s diseases (AD), Multiple Sclerosis (MS), Parkinson’s diseases (PD), and many others. While neuroinflammation is a key target for a range of neuropathological diseases, there is a lack of effective countermeasures to tackle it, and existing experimental therapies require fairly invasive intracerebral and intrathecal delivery due to difficulty associated with the therapeutic crossover between the blood-brain barrier (BBB), making such treatments impractical to treat neuroinflammation long-term. Here, we present the development of an optimal neurotherapeutic using our Nanoligomer™ discovery engine, by screening downregulation of several proinflammatory cytokines (e.g., Interleukin-1β or IL-1β, tumor necrosis factor-alpha or TNF-α, TNF receptor 1 or TNFR1, Interleukin 6 or IL-6), inflammasomes (e.g., NLRP1), key transcription factors (e.g., nuclear factor kappa-B or NF-κβ) and their combinations, as upstream regulators and canonical pathway targets, to identify and validate the best-in-class treatment. Using our high-throughput drug discovery, target validation, and lead molecule identification via a bioinformatics and AI-based ranking method to design sequence-specific peptide molecules to up-or down-regulate gene expression of the targeted gene at will, we used our discovery engine to perturb and identify most effective upstream regulators and canonical pathways for therapeutic intervention to reverse neuroinflammation. The lead neurotherapeutic was a combination of Nanoligomers™ targeted to NF-κβ (SB.201.17D.8_ NF-κβ1) and TNFR1 (SB.201.18D.6_TNFR1), which were identified using *in vitro* cell-based screening in donor-derived human astrocytes, and further validated *in vivo* using a mouse model of lipopolysaccharide (LPS)-induced neuroinflammation. The combination treatment SB_NI_111 was delivered without any special formulation using a simple intraperitoneal injection of low-dose (5mg/kg) and was found to significantly suppress the expression of LPS-induced neuroinflammation in mouse hippocampus. These results point to the broader applicability of this approach towards the development of therapies for chronic neuroinflammation-linked neurodegenerative diseases, sleep countermeasures, and others, and the potential for further investigation of the lead neurotherapeutic molecule as reversible gene therapy.

## INTRODUCTION

Although acute neuroinflammation plays a protective role in the body,^1–3^ chronic neuroinflammation is always considered detrimental and damaging to the nervous tissue due to increased levels of microgliosis and astrocytosis, oxidative stress, protein misfolding, and neurodegeneration.^4–6^ Growing evidence suggests that chronic neuroinflammation and the neurotoxic cytokine milieu associated with this persistent neuroinflammation contribute to the development of neurodegenerative diseases like AD, PD, MS, etc., and cognitive impairment.^7–12^ Innate immune activation in astrocytes and glial cells appears to be particularly important in this context,^13–16^ and recent data demonstrate a specific accumulation of pro-inflammatory “disease-associated astrocytes” in the patient’s brain.^17–19^ Further, new evidence suggests that the compounded effect of misfolded proteins leads to further glial and astrocyte activation attributed to the cytokine-like effect of amyloid-beta (Aβ)^20–22^ and immune activation in response to Aβ.^7,23,24^ Although extreme clinical interventions can include anti-inflammatory steroids, their effectiveness as neurotherapeutic countermeasures have not yet been established physiologically.^25,26^ Therefore, chronic neuroinflammation has been identified as an important neurotherapeutic target for AD, MS, PD, ischemic strokes, and others,^27^ along with other conditions such as sleep deprivation and impairment of memory and learning, especially hippocampal cognitive deficits.^28–31^

There are currently no known effective neuroinflammation countermeasures available to reduce the risk of long-term dementia, and neurological diseases. While several targets have been identified in the literature, ranging from pro-inflammatory cytokines, and inflammasomes, to transcription factors, there is no rational way to screen and rank these targets to create best-in-class and first-in-class neurotherapeutics. Overexpression of pro-inflammatory mediators such as IL-1β, IL-6, or TNF-α in the brain indicates a greater proportion of activated microglia, and hence likely a greater degree of neuroinflammation.^32,33^ Several pro-inflammatory cytokines have been implicated in multiple potential pathways such as increasing BBB permeability^34–37^ and neuroinflammation,^38^ leading to increased astrocytosis and microgliosis and neurodegenerative diseases.^37,39,40^ IL-6 plays a critical role in neuroinflammation and blocking its signaling with a monoclonal antibody against its receptor (Tocilizumab) has been approved for the treatment of several inflammatory diseases.^32,33,41^ IL-1β is another key mediator that drives neuroinflammation,^42,43^ and its inhibitor in the form of a recombinant protein is approved for treating rheumatoid arthritis and is in clinical trials for treating other chronic inflammatory diseases.^44,45^ TNFα is an important therapeutic target for developing therapies for controlling inflammation.^46^ Several recombinant proteins and antibodies that bind to and neutralize TNFα have been investigated for various inflammatory diseases. TNF-α activates two types of receptors-TNFR1 and TNFR2. TNFR1 promotes apoptosis and activates NF-κB, which leads to proinflammatory cytokine production and glial activation, and subsequent neuroinflammation and neuronal death.^47,48^ Together, IL-1 β and TNF-α have been shown to cause neuronal death synergistically by the direct effects of these cytokines on neurons or indirectly by glial production of neurotoxic substances.^49^ In addition, they induce the death of astrocytes which results in the transient production of IL-6 and other inflammatory mediators.^50^ NLR-family pyrin domain-containing 1 (NLRP1) inflammasome in the CNS is primarily expressed by pyramidal neurons and oligodendrocytes, and is considered a predominant element in the inflammatory process.^51^ NLRP1 is activated in response to amyloid-β (Aβ) aggregates, and its activation triggers a cascade of events ultimately resulting in apoptosis and axonal degeneration. Studies in murine AD models have linked upregulation of NLRP1 with neuronal death and cognitive decline.^51^ Among targeted transcription factors, NF-κβ is a central regulator of inflammation, and identified as a target in several neurodegenerative conditions like MS, AD, etc.^52–54^ Specifically, the causal link of NF-κβ activation with pathogenesis in neurodegenerative diseases through astrocytosis and microgliosis, it’s role in synaptic plasticity and depression of synaptic transmission, makes it an important potential target for countermeasure evaluation.^55^ To identify potential inflammasome targets, NLRP1 inhibition using small molecules has been reported to hinder the formation of the inflammasome complex and ATP binding for developing anti-inflammatory therapeutics.^56^

Despite remarkable progress in the field of nucleic acid therapeutics based on microRNAs, small interfering RNAs, long non-coding RNAs, and deactivated CRISPR-Cas9 technology in the recent years, several challenges associated with their delivery, stability, internalization, and target specificity need to be addressed for their successful clinical translation and development as neuroinflammation countermeasures.^57–59^ For example, technologies like deactivated CRISPR-Cas9 require tedious cloning and optimization that are time-consuming and expensive. Sachi has developed Nanoligomer™ neurotherapeutic discovery engine, a high-precision tool that generates sequence-specific, nano-biohybrid molecules called Nanoligomers™ to address the shortcomings of the existing methodologies.^60^ A Nanoligomer has six design elements with peptide nucleic acid (PNA), a synthetic DNA-analog where the phosphodiester bond is replaced with 2-N-aminoethylglycine units, as the nucleic acid binding domain^61^ and can be designed to either up-or down-regulate any desired gene by either binding to its mRNA or DNA.^60^ Nanoligomers offer improved stability, facile delivery and internalization, high target specificity (on-target) and minimal off-targeting, and can be easily designed to rapidly create a library of neurotherapeutic molecules with the capability to cross the BB using any route of administration (intraperitoneal IP, intranasal IN, or intravenous IV), superior biodistribution and minimal accumulation compared to existing nucleic acid therapeutics.^59^ In addition, they modulate related gene network targets ultimately resulting in pathway modulation. Previously, we have demonstrated that Nanoligomer technology is very effective in identifying immunotherapy lead targets and molecules for countermeasure development.^60^

To screen and rank these targets and identify the best-in-class and potential first-in-class therapeutic molecule, we applied Sachi’s Nanoligomer discovery engine to regulate the above-mentioned neuroinflammation relevant targets (pro-inflammatory cytokines, inflammasome, transcription factors, and combinations) to identify the most effective upstream regulator for developing countermeasures for chronic neuroinflammation.

## RESULTS AND DISCUSSION

Nanoligomer™, a nano-biohybrid molecule enables both up-and down-regulation of any gene of interest. Peptide nucleic acid (PNA), a synthetic DNA-analog in which phosphodiester bond is replaced with 2-N-aminoethylglycine units,^61^ is the nucleic acid-binding domain of the Nanoligomer™. PNAs afford stronger hybridization, target specificity,^62^ and higher stability in human blood serum and mammalian cellular extracts^63^ compared to naturally occurring RNA or DNA. Nanoligomer downregulators act either *via* targeting of mRNA to inhibit RNA stability or protein expression or *via* targeting of DNA.^59,60^ Nanoligomer upregulators trigger transcriptional activation by binding at the respective promoter region and attachment of specific domains that recruit transcriptional activators (Fig 1).

**Fig.1.**
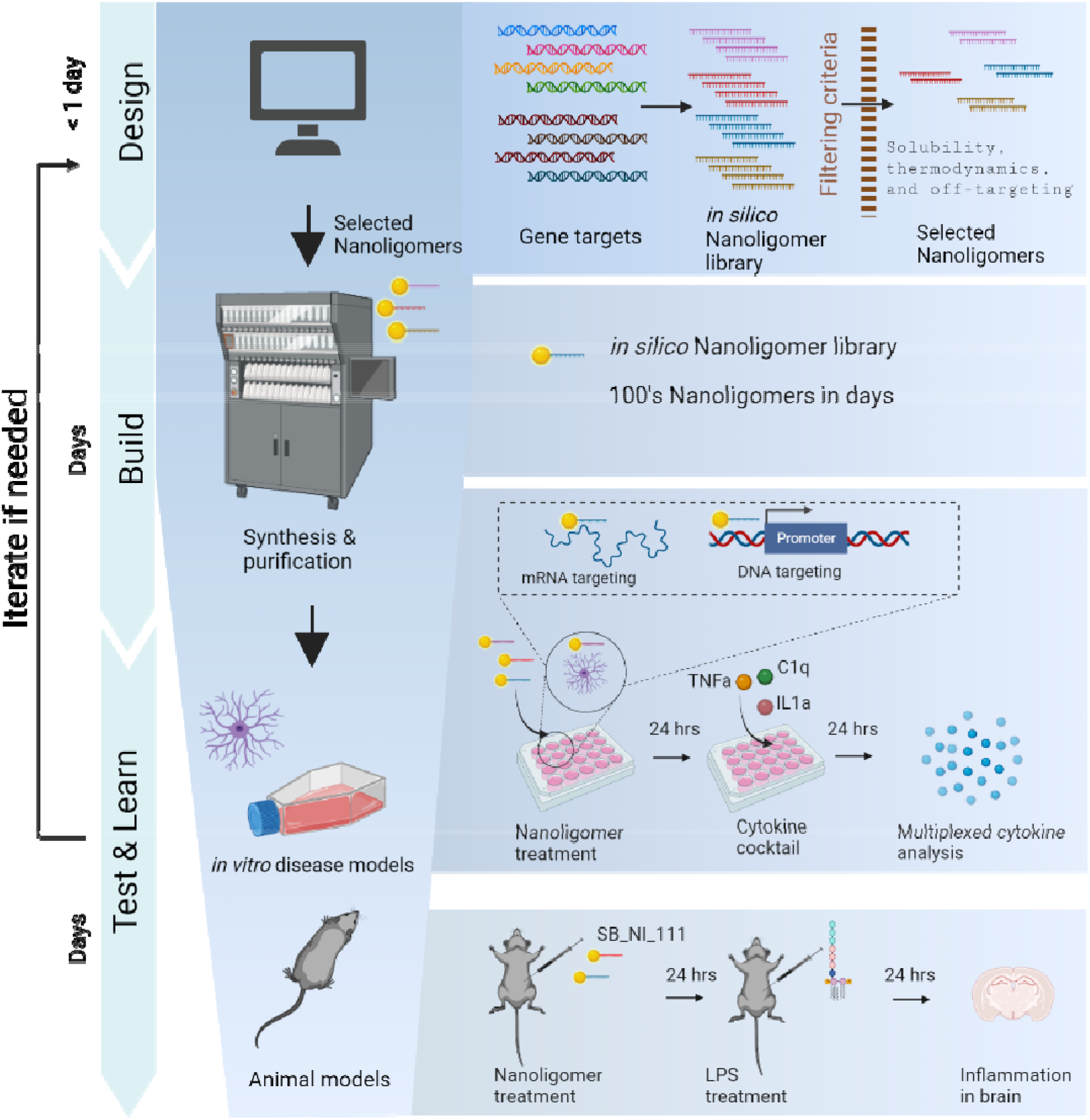
Nanoligomer-based discovery engine for neurotherapeutic discovery. (A) Process flow for the Nanoligomer platform follows four stages Design, Build, Test, and Learn. (Top) An in silico Nanoligomer library is created for each selected gene target and all candidates are evaluated and selected through a four-step process. First, gene target regions (RNA and/or DNA) are identified based on factors such as sequence composition and predicted transcription or translational regulatory regions. These seed regions are then evaluated for expected synthesis and solubility considerations followed by filtering based on undesirable features such as self-complementary and direct match or multiple mismatch off-targets. The fourth step is a thermodynamic analysis using a Naïve Bayes classifier for target vs off-target binding and optimization of possible on-target effectiveness. (Middle Left) Best Nanoligomer designs are synthesized, purified, and screened in astrocytes via a multiplex cytokine panel. (Middle Right) Inset shows the two mechanisms of action by which Nanoligomers can regulate gene expression. Left: Nanoligomers target mRNA to regulate RNA stability or translation. Right: Nanoligomers target genomic DNA to regulate transcription. Nanoligomers target genomic DNA to regulate transcription. (Bottom) Finally, an identified Lead Nanoligomer candidate is tested in a murine model of LPS-induced neuroinflammation and evaluated by a cytokine panel for its ability to reduce neuroinflammation.

### Nanoligomer™ Discovery Engine Workflow

Sachi’s Nanoligomer™ Discovery Engine consists of four stages, namely, Design, Build, Test, and Learn (Fig 1). The Nanoligomer Design stage utilizes Sachi’s bioinformatics toolbox in which the sequences of the identified gene targets are used as inputs to create an *in silico* Nanoligomer library for each selected gene target (Fig 1). Next, all the candidates were evaluated to select top Nanoligomers using the following four steps: 1) identification of the gene target regions (RNA and/or DNA) based on sequence composition and predicted transcription or translational regulatory regions; 2) evaluation of the seed regions for expected synthesis and solubility considerations; 3) filtering based on undesirable features such as self-complementarity and direct match or multiple mismatches (up to two) off-targeting effects for each candidate across the human genome; and, 4) thermodynamic analysis using a machine-learning approach based on Naïve-Bayes classifiers^64–68^ for target vs off-target binding and optimization of possible on-target effectiveness. These steps were iterated to design Nanoligomers targeting different sites (on the DNA or mRNA) of the target of interest. Next, all the candidates in the bioinformatics generated *in silico* Nanoligomer library were scored and ranked as described in detail elsewhere.^60^ Following this, top Nanoligomer designs (at least three per target-one per genomic location) were synthesized as a single modality using high throughput peptide synthesizer and purified (Fig 1, Table S1). A Missense Nanoligomer (designed with no homology to any gene in the specific genome at the DNA/ mRNA level) was also designed and synthesized to rule out any possible non-specific effects. Top Nanoligomers were then screened in an *in vitro* model of neuroinflammation (human astrocytes treated with cytokine cocktail) using multiplex cytokine panel to identify Lead Nanoligomer for each target (Fig 1). Finally, *in vitro* validated Lead Nanoligomer candidate(s) was evaluated in a murine model of neuroinflammation for its ability to reduce or alleviate neuroinflammation.

### Target and Nanoligomer Ranking Based on *in vitro* Screening

The selected neuroinflammation-relevant targets, viz., NLRP1, IL-6, IL-1β, TNF-α, TNFR1, and NF-κβ and their corresponding Nanoligomers (at least three per target, for TNFR1-splice variants 1 and 2 were used) were ranked based on screening in an *in vitro* human astrocyte-based model of neuroinflammation and multiplex cytokine analysis (Fig 1; Fig 2). Astrocytes are considered instrumental in maintaining brain homeostasis, the blood-brain barrier (BBB), and regulating inflammatory responses in the central nervous system.^69^ They can attain either neurotoxic (A1-phenotype) or neuroprotective (A2-phenotype) state based on the surrounding milieu and stimuli. Changes in the relative population of A1 and A2 phenotypes (more A1 and less A2, respectively) have been observed in neuroinflammation and neurodegenerative diseases.^69^

**Fig 2.**
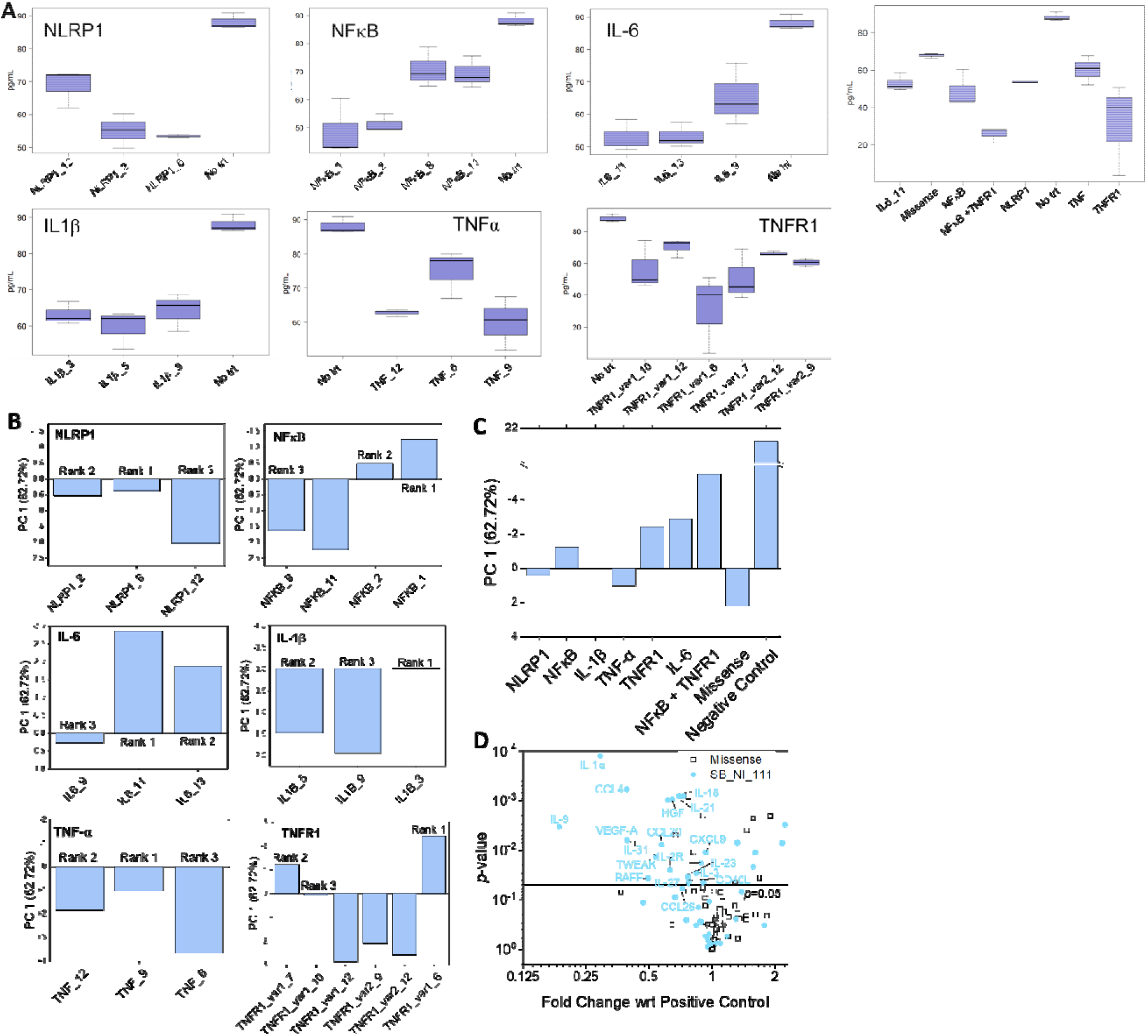
Lead molecule and target identification from high-dimensional datasets using principal component analysis (PCA). (A) Cytokine-based sorting using IL-1α (in pg/ml) used to identify the best Nanoligomer candidate for each target, and then ranking of respective targets. PCA of multiplexed ELISA data for activated astrocytes cytokine expression data for various Nanoligomer treatments and untreated samples. (B) PC1 data for multiple Nanoligomer designs targeting NLRP1, NFκB1, IL-6, IL-1β, TNFα, and TNFR1. (C) PC1 data for best Nanoligomer designs for NLRP1, NFκB1, IL-6, IL-1β, TNF-α, and TNFR1 targets along with missense and no Nanoligomer/ no cytokine treatment controls. PCA is based on cytokine data obtained from three biological replicates. (D) Volcano plot of high-dimensional 65-plex cytokine data normalized with respect to cytokine only treatment (positive control) shows that the cytokines were upregulated due to cytokine treatment (neuroinflammatory state), were restored back to the lower healthy levels by the SB_NI_111 Nanoligomer treatment. Also, the missense Nanoligomer control shows values between the scrambled sequence and cytokine only treatment are very similar (clustered around 1), indicating the effect observed here is due to sequence-specific action of the Nanoligomer treatment.

A human astrocyte-based *in vitro* model of neuroinflammation was created using the cytokine cocktail (TNF-α, IL-1α, and C1q) known to induce astrocytes toward neuroinflammatory A1-phenotype.^48,70^ Briefly, astrocytes were first pre-treated with the gene-specific 10 μM Nanoligomers (with or without cytokine cocktail) for 24 hours, after another 24 hours media supernatants were collected and analyzed for secreted protein/cytokine expression (see methods for details). First, we used IL-1α expression level as a key proinflammatory biomarker (Fig. 2A), to rank the respective targets. Next, the high-dimensional (65-plex) cytokine data was normalized to their respective cytokine levels in non-treated astrocytes and significance was calculated using a t-test between replicates. Any cytokines that showed broad expression changes across multiple treatments were removed from analysis to elucidate gene-specific effects. Next, the data was transformed using principal component analysis (PCA) to reduce dimensionality to find trends and patterns as described in detail elsewhere.^60,71^ Based on the Scree plot (Fig. S1), Principal Component 1 (PC1) captured most of the variance (62.72%) in the data. PC2, PC3, and PC4 captured only 10.79%, 6.87%, and 4.33% of variability, respectively, suggesting that PC1 scores alone could be used for target/ Nanoligomer ranking. Based on the PC1 scores (higher negative values imply better efficacy of Nanoligomer downregulator), we first ranked the various Nanoligomer designs for each gene target (Fig. 2B) and then selected the top-ranked (Rank 1) Nanoligomer downregulator for each target for ranking the six targets and one combination investigated in this study. Both positive (Missense Nanoligomer, represents diseased state with no treatment of a Nanoligomer but with cytokine cocktail-induced neuroinflammation) and negative control (no treatment no cytokine control, represents normal healthy state, no neuroinflammation) were used. (Fig. 2C).

A more detailed examination of the 65-plex cytokine data using volcano plot, normalized with respect to cytokine treatment with no Nanoligomer treatment (positive control) revealed that: 1) Missense Nanoligomer (scrambled sequence with no overlap) data is similar to no treatment (most cytokines are close to 1, and p-values are high/not significant); and 2) Nanoligomer treatment (for the top-ranked combination) significantly reduced the inflammatory cytokines to much lower values (e.g., IL-1α) with high significance (Fig. 2D). This reveals that while such detailed, high-dimensional data would be invaluable to look at expression and impact on individual cytokines, getting a consistent ranking for several Nanoligomer targets and treatments would be infeasible. Therefore, we reduced the dimensionality of the data using PCA, to rank different targets used in the treatments.

Based on IL-1α expression level (Fig. 2A), PCA analysis of multiplexed 65-plex cytokine data (Fig. 2B-D), heat maps, and unsupervised hierarchical and K-means clustering for astrocyte and PBMCs for different targets (Fig. S2), we made the following assessments: 1) NLRP1_6, NF-κβ _1, IL-1B_3, TNF_9, TNFR1_var1_6, and IL6_11 were the best Nanoligomers for their respective targets. 2) Target Ranking: NF-kB and TNFR1 combination (Rank 1), IL-6 (Rank 2), TNFR1 (Rank 3), NF-κβ (Rank 4), IL-1 β (Rank 5), NLRP1 (Rank 6), and TNF-α (Rank 7). The PCA and ranking analysis suggested IL-6 (Rank 2) as a promising single gene target for neuroinflammation (based on principal component values farthest from the missense control and closest to the negative control). However, IL-6 has a broad role in mediating several key physiological pathways and fulfills multiple contrasting functions: 1) essential homeostatic functions, which include immune cell proliferation and differentiation as well as metabolic functions, and 2) pro-inflammatory actions due to dysregulated activity.^72^ In the past, monoclonal antibodies against IL-6 or IL-6 receptor (IL-6R) and Janus kinases (JAK) inhibitors showed some promise in clinical trials, but their use resulted in compromised immune defense and exposing the host to a range of debilitating infections and diseases.^72^ Therefore, we selected the top-ranked combination therapy for neuroinflammation treatment. Combination or “cocktail” therapy is an accepted paradigm for a variety of complex diseases.^73^ Drug interaction means how another drug affects the activity of the selected drug when both are administered together. This action can be synergistic (when the drug’s effect is increased) or antagonistic (when the drug’s effect is decreased).^74^ We used Bliss Independence model^75^ to quantify synergy and combinatorial effects. Synergy (*S*) factor is the deviation from no interaction and is defined as: *S* = PC1_AB_ - (PC1_A_ x PC1_B_), where PC1_A_, PC1_B_, and PC1_AB_, are the PC1 values for the target (and Nanoligomer) A, B, their combination AB, respectively. *S* >□0 means synergy and *S* <□0 means antagonism. Based on this analysis, the combination of TNFR1 and NF-κβ targets and their respective Nanoligomers emerged more effective (*S* factor = 2.53) as compared to individual targets and Nanoligomers (Fig 2B). Therefore, we decided to further validate combinatorial therapy with TNFR1 (Sachi’s Library Name: SB.201.18D.6_TNFR1) and NF-κβ (Sachi’s Library Name: SB.201.17D.8_ NF-κβ1), named SB_NI_111, in an *in vivo* setting.

### A low dose of SB_NI_111 reverses LPS-induced neuroinflammation in a mouse model

To evaluate combinatorial or “cocktail” therapy with TNFR1 and NF-κβ *in vivo*, we first designed and synthesized Nanoligomers targeting mouse TNFR1 and NF-kB genes using Sachi’s Nanoligomer pipeline (as described earlier). TNFR1 (Sachi’s Library Name: SB.24D.8_TNFR1) and NF-κβ (Sachi’s Library Name: SB.25D.5_NFKB1) Nanoligomers were mixed in equal amounts (mg) to make Nanoligomer cocktail SB_NI_111. By examining the serum cytokines involved in several pathological immune responses, we previously established the non-immunogenicity and non-toxicity of Nanoligomers *in vivo*.^60^ In addition, we have shown that Nanoligomer can be efficiently delivered *in vivo* through the peritoneum and have low dissociation constant (K_D_), facilitating excellent therapeutic action even at low doses. IP injection of GM-CSF downregulator showed Nanoligomer in the brain within 1 hour of injection suggesting crossing the BBB.^59^ We used Lipopolysaccharide (LPS), which has been used widely *in vitro* and *in vivo*, to model neuroinflammation associated with neurodegeneration.^6^ LPS has been known to activate various intracellular molecules *via* receptor-mediated binding and alter the expression of a plethora of inflammatory mediators, ultimately initiating the development of neurodegenerative processes. Previous studies have shown that the intraperitoneal (i.p.) injection of LPS increases levels of pro-inflammatory TNF-α, IL-1α, IL-1β, and IL-6 in the plasma and brain regions (hippocampus and cortex).^76,77^ In addition, pro-inflammatory cytokines, such as IL-1β, IL-6, and TNF-α, produced by a systemic inflammatory response can also reach the CNS (hippocampus and cortex) through the blood circulation,^78^ suggesting that LPS administration is ideal for mimicking neuroinflammatory state and associated neurodegeneration and neuropathological pathways.^6^

Next, we evaluated SB_NI_111 (24D.8_TNFR1 (5mg/Kg) and 25D.5_NFκB1 (5mg/Kg)) cocktail in *vivo*. The mice were split into three treatment groups (3 mice/ group), which were administered either vehicle (PBS) negative control (Group 1 and 2) or SB_NI_111 cocktail (Group 3) *via* intraperitoneal (i.p.) injection (Day −1). Twenty-four hours post-injection (hpi), Group 1 (saline, negative control) mice were injected with saline, and Group 2 (positive control, represents neuroinflammatory state) and group 3 (SB_NI_111, Nanoligomer treated) were injected with 0.75mg/Kg of LPS (Day 0) (Fig 3A). Six hours after that, mice were euthanized by carbon dioxide, brains were dissected, and hippocampus tissues were harvested, flash-frozen, and stored at −80 °C until further use. Throughout the study, no signs of distress were observed in mice. Frozen mouse tissues were processed and analyzed for cytokines and chemokines analysis using 36-Plex Mouse ProcartaPlex Panel 1A (ThermoFisher Scientific, see methods for details). Additionally, gene expression measurements of targeted TNFR1 and NFKB1 genes in mouse hippocampal tissue confirmed one-log (∼2-fold reduction) for each of the respective targets, compared to the no treatment sample (Fig. S3B,C). The high-dimensional cytokine data (normalized with respect to negative control) was analyzed using PCA, which revealed that most of the variance (59.46%) in the data was captured by principal component 1 (PC1) (Scree plot, Fig S3A). PC2 captured 21.45% variance, followed by PC3 and PC4, which captured only 8.29% and 5.34% of variability, respectively. PC1 vs PC2 plot (Fig 3B) clearly shows a distinction between SB_NI_111 (TNFR1+NFκB) cocktail treated and untreated LPS-exposed mice, establishing that SB_NI_111 treatment results in broad alterations in the cytokine profile in mouse hippocampus.

**Fig 3.**
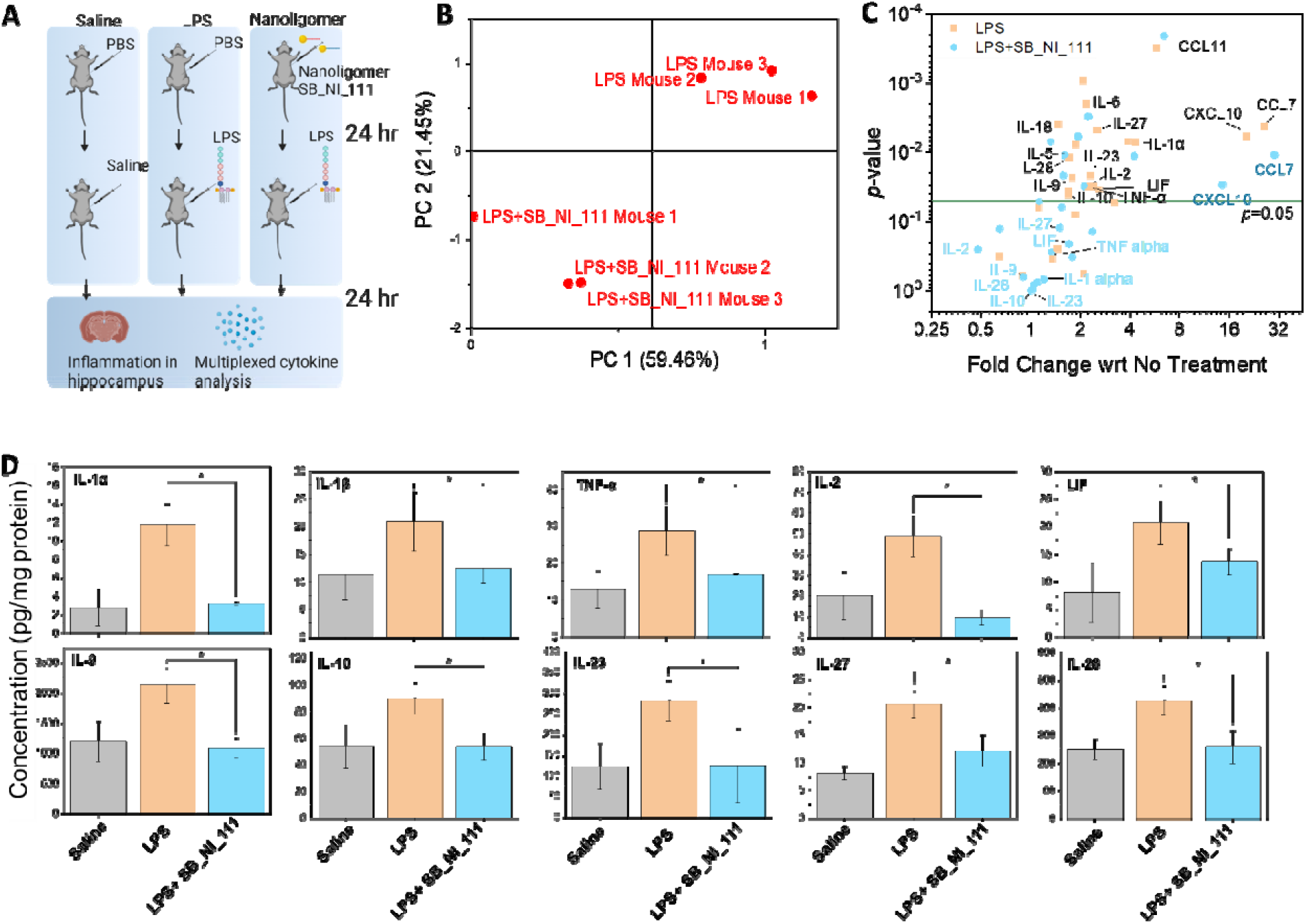
In vivo evaluation of SB_NI_111 cocktail in an LPS-induced neuroinflammation mouse model. A) Schematic showing experimental design B) Principal component analysis shows a distinction between irradiated SB_NI_111 treated and untreated LPS-induced mice. C) Volcano plot and D) bar plots show that the cytokines (IL-1α, IL-1β, TNFα, IL-2, LIF, IL-9, IL-10, IL-23, IL-27, and IL-28) that were upregulated due to LPS treatment (neuroinflammatory state), were restored back to the normal healthy levels (saline, negative control) (*p<0.05) upon treatment with the SB_NI_111 cocktail. Data shown is an average of three biological replicates, and error bars represent standard deviation. The statistical significance was determined using unpaired two-tailed Students *t*-test.

LPS administration significantly upregulated the expression of several cytokines (w.r.t. vehicle, saline), viz, IL-1α, IL-1β, TNFα, IL-2, LIF, IL-9, IL-10, IL-23, IL-27, IL-28, and (Fig 3C, D). IL-1α, IL-1β, and TNF-α are considered the main mediators of neuroinflammation, and their expression levels were significantly increased as a result of LPS treatment, mimicking a neuroinflammatory state. Treatment with the SB_NI_111 cocktail significantly downregulated the expression of all these cytokines and restored the cytokines levels either back to or even lower than the basal level (saline, negative control, no neuroinflammation) (p<0.05) (Fig. 3C, D). In addition, levels of IL-2, LIF, IL-9, IL-10, IL-23, IL-27, and IL-28 were also restored back to no disease state (saline, negative control) (Fig 3 C, D). However, some chemokines, notably CXCL10, CCL7 and CCL11 were significantly higher, compared to no treatment, which is not unexpected given their gene expression is largely unaffected by targeted genes. However, some other cytokines like IL-5, IL-18, and IL-6 were also not fully restored to pre-inflammation levels, likely due to extremely high inflammatory response of LPS. These results are especially significant since LPS administration and establishment of the associated behavioral data has shown that this neuroinflammation model is directly linked to hippocampal learning deficit and cognitive decline, along with the clear causal onset of neurodegenerative diseases, as evidenced by increased Aβ load neuronal death.^6^ Using this well-established model and identified biomarkers, the reversal of all key mediators like IL-1α, IL-1β, and TNF-α to the original state (saline levels, Fig. 3D) and the similarity of the high-dimensional dataset between saline and the LPS model with the lead Nanoligomer treatment (Fig. 3C) validates the *in vivo* efficacy and potential advancement and assessment of the developed countermeasure for further translation. These results establish the efficacy of Sachi’s SB_NI_111 cocktail in significantly suppressing the neuroinflammation in the hippocampus of the mouse model. Future studies with SB_NI_111 cocktail will explore variable dosing regimens, an alternate route of administration such as intranasal, as well as multiplexing with other pathway targets for controlled mitigation of neuroinflammation long term.

## CONCLUSIONS

In this study, we demonstrated the applicability of Sachi’s Nanoligomer™ discovery engine for high-throughput and rapid neurotherapeutic development to counter chronic neuroinflammation, as a neurotherapeutic target for AD, MS, PD, ischemic strokes, and as a countermeasure for sleep-deprivation and impairment of memory and learning, especially hippocampal cognitive deficits. Our discovery tool utilized precise regulation and gene perturbation of a number of upstream immune regulators and canonical pathways up and down using donor-derived human astrocytes, and multi-dimensional PCA analysis to identify the combination of the most efficacious target (NF-κβ and TNF-R1 combination termed as SB_NI_111). Then, we demonstrated that LPS-induced neuroinflammation in a mouse model can be reversed through precise targeting of NF-κβ and TNF-R1 *via* SB_NI_111 cocktail in the mouse brain. The Nanoligomer-induced reversible gene expression modulation can have important therapeutic applications by combining specificity and ease of delivery, with the ability to target key gene networks to alleviate dysfunction in a number of key protein expression. Sachi’s Nanoligomer™ Discovery Engine represents both a strong key target ID platform, as well as a therapeutic modality.

## MATERIALS AND METHODS

### Nanoligomer Design and Synthesis

DNA and transcript sequences of gene targets of interest were used as input to Sachi’s bioinformatics toolbox. The gene sequences/ transcripts were interrogated for optimal design based on sequence identity, predicted gene regulatory sites, thermodynamic binding optimization, self-complements, off-target effects, solubility, and synthesis parameters.^59,60^ Further, machine learning based on Naïve-Bayes classifiers and experimental data was used to rank, and then validate, the different candidates for gene-perturbation of the potential target.^64–68^ This analysis was used to select the Nanoligomer candidates with the least off-target effects. Following design and ranking, a Nanoligomer was selected and synthesized (Table S1) using our high throughput, automated peptide synthesizer, AAPPTEC Vantage (AAPPTEC, LLC) with solid-phase Fmoc chemistry at a 10-μmol scale on 4-methylbenzhydrylamine (MBHA) rink amide resin. Fmoc–Nanoligomers monomers were obtained from PolyOrg Inc., with A, C, and G monomers protected with Bhoc groups. Purified Nanoligomers’ were quantified by UV-Vis (Thermo Scientific NanoDrop, at 260 and 400 nm, respective extinction coefficients for SB_NI_111 are ε_260_= 213,800 cm^-1^; ε_400_= 10,000 cm^-1^) according to known optical parameters and then stored at 4 °C until further use.

### Astrocyte culture and treatments with Nanoligomers

Primary human astrocytes (Lonza) were maintained in a complete astrocyte growth medium (ScienCell) at 37 °C and 5% CO2 in a humidified incubator and subcultured at ∼80-90% confluency. Stock solutions of the cytokines (TNF-α, IL-1α, C1q-Sigma-Aldrich) were prepared in molecular biology grade water, aliquoted, and stored at −80ºC. The composition of the cytokine cocktail used for astrocyte activation was: TNF-α: 30 ng/mL, IL-1α: 3 ng/mL, and C1q: 400 ng/mL (effective concentrations in the culture medium).^48,70^ For Nanoligomer treatment, 10,000 cells were seeded per well of a 48-well plate and cultured until 80% confluency. Then, astrocytes were first pre-treated with the gene-specific Nanoligomers (with or without cytokine cocktail) for 24 hours, after another 24 hours media supernatants were collected and analyzed for secreted protein/cytokine expression.

### Quantification of secreted proteins using Immune Monitoring 65-Plex Human ProcartaPlex™ Panel

Cell culture supernatants were analyzed for secreted proteins using Immune Monitoring 65-Plex Human ProcartaPlex™ Panel for MAGPIX (Thermofisher Scientific, Carlsbad, CA) as per the manufacturer’s instructions. This is a preconfigured multiplex immunoassay kit that measures 65 cytokines, chemokines, and growth factors for efficient immune response profiling, biomarker discovery, and validation. It consists of two separate kits: a 43-plex and a 22-plex kit that has been specifically designed to be used on the MAGPIX instrument that can detect up to 50 targets simultaneously. In this study, astrocyes supernatants were probed for the following 65 markers: APRIL, BAFF, BLC, CD30, CD40L, ENA-78, Eotaxin, Eotaxin-2, Eotaxin-3, FGF-2, Fractalkine, G-CSF, GM-CSF, GROα, HGF, IFN-α, IFN-□, IL-1α, IL-1β, IL-2, IL-2R, IL-3, IL-4, IL-5, IL-6, IL-7, IL-8, IL-9, IL-10, IL-12p70, IL-13, IL-15, IL-16, IL-17A, IL-18, IL-20, IL-21, IL-22, IL-23, IL-27, IL-31, IP-10, I-TAC, LIF, MCP-1, MCP-2, MCP-3, M-CSF, MDC, MIF, MIG, MIP-1α, MIP-1β, MIP-3α, MMP-1, NGF-β, SCF, SDF-1α, TNF-α, TNF-β, TNF-R2, TRAIL, TSLP, TWEAK, and VEGF-A. The plate was read using the MAGPIX instrument and xPONENT® software (version 4.2, Luminex Corp, Austin, Texas, US). The overall intra- and inter-assay precision are reported by the manufacturer as 2–19%, and accuracy as 87–107% over the calibration range of 3.2–10,000□pg/mL protein concentration. Data analysis was performed in Microsoft Excel and plots were made using OriginLab software.

### Murine model study

The animal care facilities at the University of Colorado at Boulder are AAALAC accredited, and all animal work was approved by the Institutional Animal Care and Use Committee at the University of Colorado. Male C57BL/6 mice (Charles River Laboratories) between 3 and 5 months old were divided into three groups. On day 0 No Treatment naïve group was injected with sterile saline PBS, then sterile saline IP 24 hours later on day 1, then sacrificed on day 2 (after 24 hours, Fig. 3A). Similarly, the sham treatment group and Nanoligomer treatment groups were injected with sterile PBS (sham treatment) and 5 mg/kg Sachi molecules respectively on day 0, then 0.75 mg/kg LPS IP on day 1 (after 24 hours), then sacrificed at on day 2 (total 48 hours). The mice were weighed the morning of each injection to determine proper dosing, and were allowed to habituate for 1 hour prior to dissection. Subjects were given a 180 mg/kg injection of Euthasol (Virbac 710101) as an anesthetic agent. Anesthetic depth was checked via toe pinch, then mice were perfused with PBS to remove blood from the brain. Hippocampus were dissected and flash frozen on dry ice and stored at −80C until homogenization. LPS was prepared by diluting in water to a final concentration of 5mg/mL. All samples were immediately flash frozen and stored at −80°C until further use.

### Quantification of Cytokine & Chemokine in mouse tissues

Flash-frozen tissues were homogenized using a mortar and pestle and syringe disruption to form a homogenate solution in tissue cell lysis buffer (EPX-99999-000). Samples were then centrifuged at 16,000xg for 10 min at 4°C and supernatant was transferred to a fresh tube. Bio-Rad’s DC Protein Assay Kit was used to determine protein content and all samples were diluted to 10 mg protein/mL. Quantification of cytokines/ chemokine was performed with Cytokine & Chemokine Convenience 36-Plex Mouse ProcartaPlex Panel 1A (EPXR360-26092-901), analyzed on a Luminex MAGPIX xMAP instrument, and quantified using xPONENT software. For standard curves, eight four-fold dilutions of protein standards were used. Target list includes: ENA-78 (CXCL5), Eotaxin (CCL11), GRO alpha (CXCL1), IP-10 (CXCL10), MCP-1 (CCL2), MIP-1 alpha (CCL3), MIP-1 beta (CCL4), MIP-2 alpha (CXCL2), RANTES (CCL5), G-CSF (CSF-3), GM-CSF, IFN alpha, IFN gamma, IL-1 alpha, IL-1 beta, IL-2, IL-3, IL-4, IL-5, IL-6, IL-9, IL-10, IL-12p70, IL-13, IL-15/IL-15R, IL-17A (CTLA-8), IL-18, IL-22, IL-23, IL-27, IL-28, IL-31, LIF, MCP-3 (CCL7), M-CSF, TNF alpha.

### Statistics and Data Analysis

Principal Component Analysis (PCA) was conducted using multivariate analysis using Software R and Origin. Up to 4 principal components were utilized in the analysis, assessed using Scree plots, and the most significant component(s) used in ranking the Nanoligomers. The heatmaps, unsupervised hierarchical and K-means clustering was carried out using Broad institute’s Morpheus tool. All statistical comparisons including ANOVA and unpaired two-tailed Student’s *t*-test was conducted using Microsoft Excel.

## ASSOCIATED CONTENT

### Supporting Information

Scree plots for PCA analysis, heatmaps, unsupervised hierarchical and K-means clustering, gene accession numbers and PNA sequences, and gene expression measurements for in vivo hippocampus tissue.

### Author Information

#### Corresponding Author

*

#### Author Contributions

P.N. conceived the idea and designed the experiments, and synthesized the Nanoligomers. S.S. conducted all the *in vitro* experiments and biochemical characterization. C.B., J.H., and M.A.G conducted the LPS mouse study. S.S. and P.N. conducted the ELISA measurements. P.N, S.S., and A.C. wrote the manuscript with input from all the authors. All authors read the manuscript and provided input.

#### Notes

S.S., A.C., and P.N. are employed by Sachi Bioworks where this technology was developed.

A.C. and P.N. are the founders of Sachi Bioworks, and P.N. has filed a patent on this technology.

## ACKNOWLEDGMENTS

This work was supported by financial support from National Aeronautics and Space Administration SBIR Contracts 80NSSC21C0242 and 80NSSC22CA116 to Sachi Bioworks. The authors thank Colleen Courtney for helping with mouse tissue ProcartaPlex sample preparation.

## TOC GRAPHIC

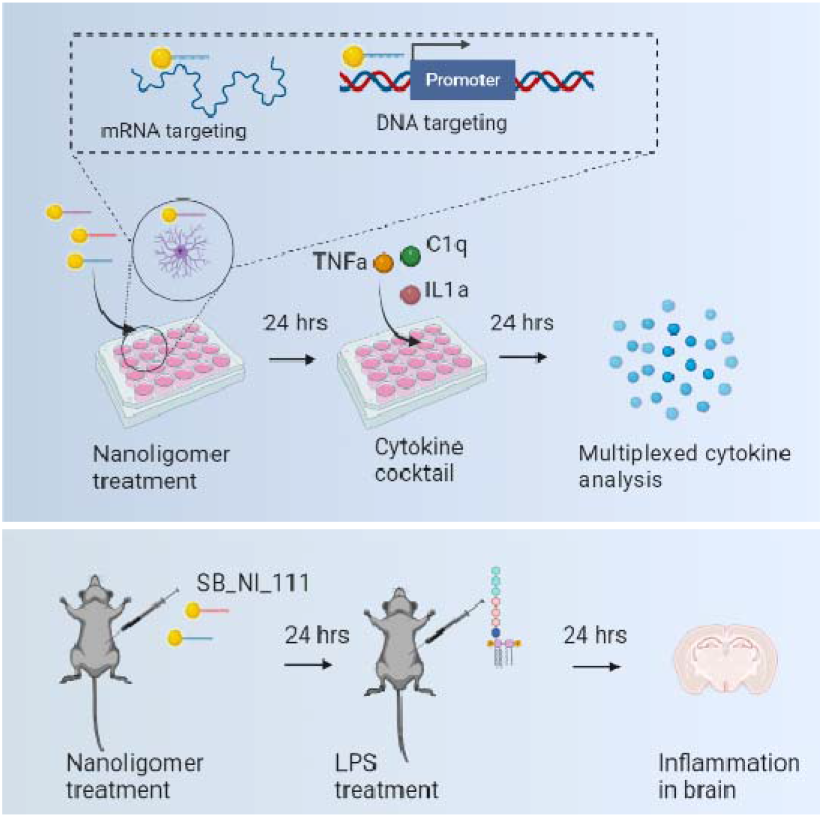

