## Supporting Information for "Identifying Optimal Neuroinflammation Treatment Using Nanoligomer™ Discovery Engine"

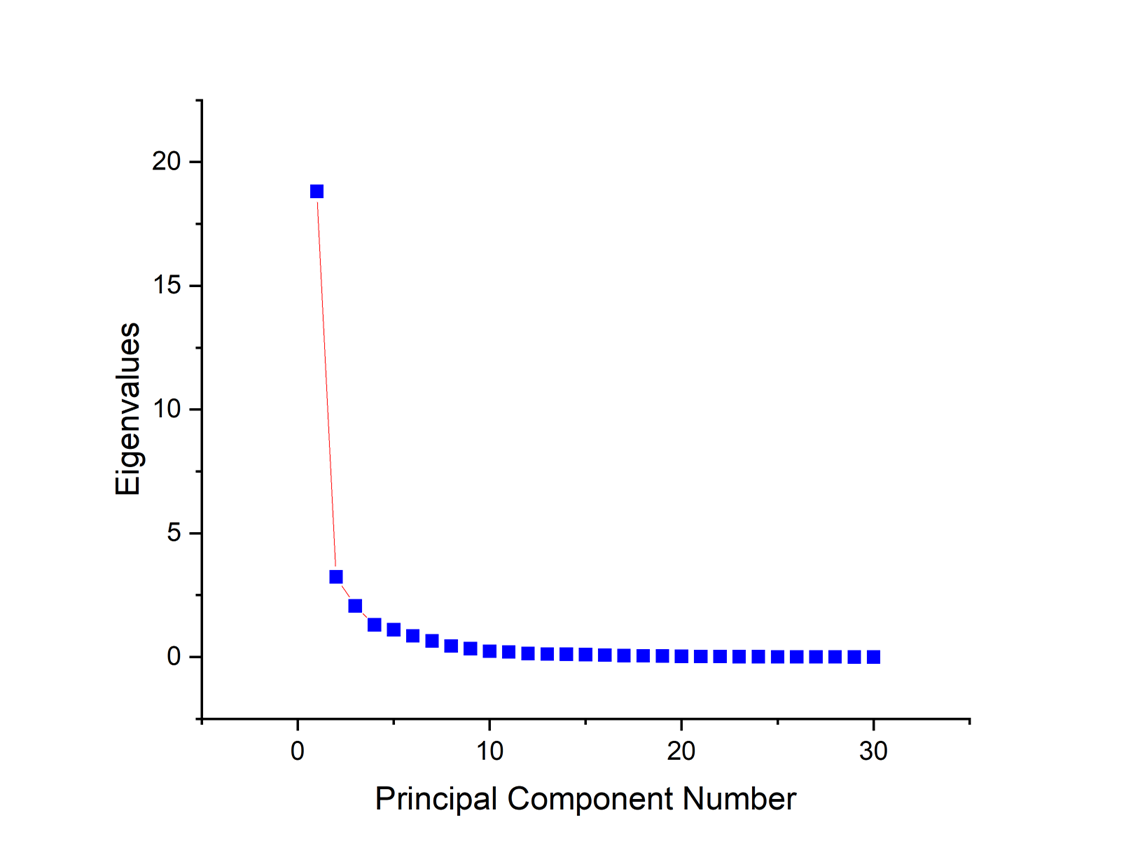

**Fig. S1** **Scree plot of principal component analysis for human astrocytes screening** using 65-plex cytokine panel. Based on this plot, Principal Component 1 (PC1) captures most of the variability (62.72%) in the data. PC2, PC3, and PC4 captured only 10.79%, 6.87%, and 4.33% of variability.

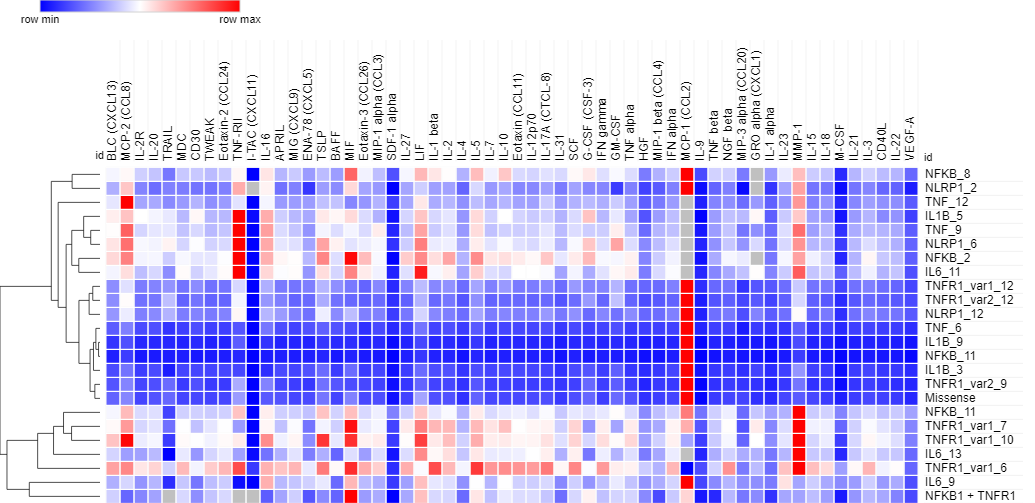

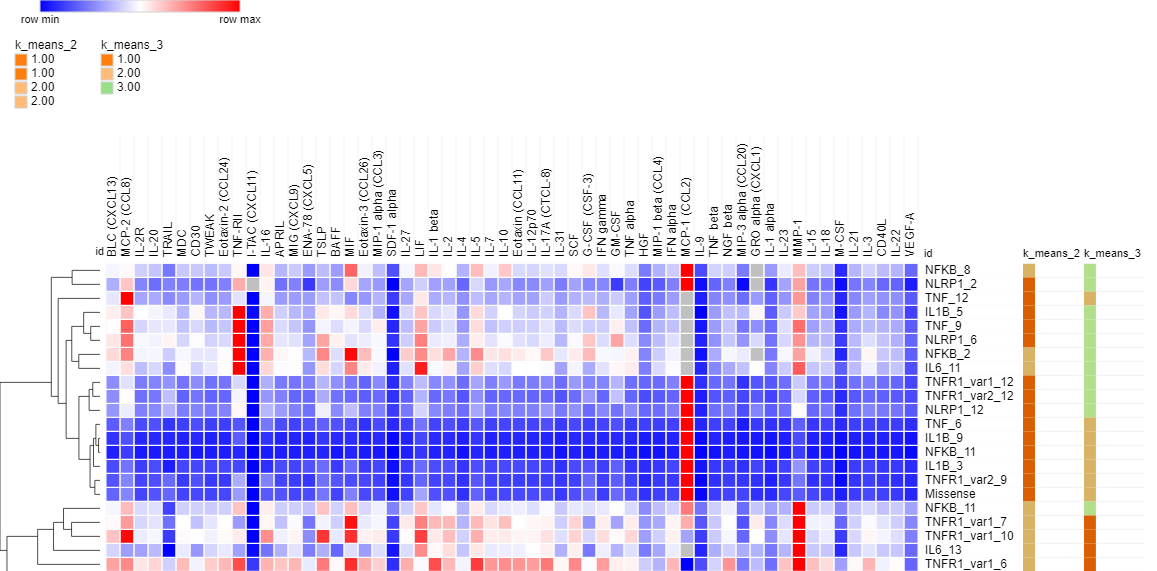

**Fig. S2**. (A) Heat maps and unsupervised hierarchical clustering for entire data set, and (B) added K-means clustering. Using the broad institute’s Morpheus tool, the heatmap and clustering support the same analysis and conclusions as the PCA analysis.

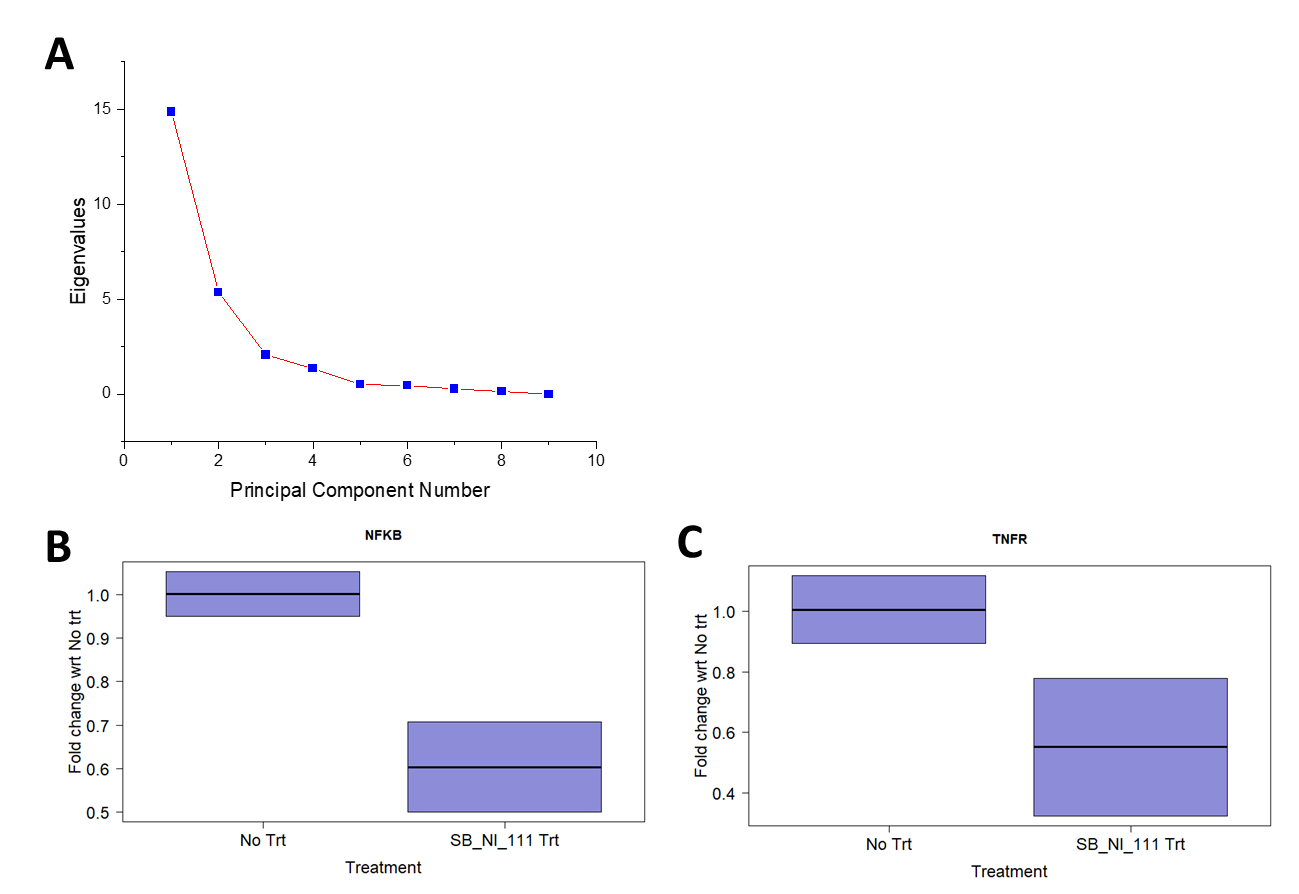

**Fig. S3** (A) Scree plot of the principal component analysis of the mouse cytokine data (LPS, LPS+SB_NI_111) using a 36-plex cytokine panel. Principal component analysis of the cytokine data suggests that principal component 1 (PC1) captures most of the variability (59.46%) in the data. PC2, PC3, and PC4 capture only 21.45%, 8.29%, and 5.34% of variability, respectively. Gene expression measurements in mouse hippocampus with respect to No Treatment. Measurements of (B) NFKB (average reduction 40.5%, p=0.05) and (C) TNFR1 (average reduction 49.74%, p=0.13), show orthogonal gene expression measurements *in vivo* in mice hippocampus. Actin B (ACTB) primers were used as housekeeping/reference gene.

**Table S1**. The PNA sequence of Nanoligomers™ shown in **Fig. 2D**.

| **Gene Accession Number** | **Gene Target** | **PNA Sequence (N to C)** |
| --- | --- | --- |
| NM_001033053.3 | NLRP1_12 | GCCATCTCTGTCCCGGA |
| NM_003998.4 | NFKB_1 | GATCATCTTCTGCCATT |
| NM_001065.4 | TNFRIvar1_6 | GAGAGGCCCATGCCAGA |
| NM_000576.3 | IL1B_3 | TACTTCTGCCATGGCTG |
| NM_000594.4 | TNF_9 | CTCATGGTGTCCTTTCC |
|  | Missense | GAGTTCATAGCTGGGCT |
| NM_000600.5 | IL6_11 | TCATAGCTGGGCTCCTG |
